# Infanticide is driven by unfamiliarity with offspring location and associated with androgenic shifts in mimic poison frogs

**DOI:** 10.1101/2024.04.11.589025

**Authors:** Amaris R. Lewis, Billie C. Goolsby, Bryan H. Juarez, Madison P. Lacey, Lauren A. O’Connell

## Abstract

Infanticide is widespread across the animal kingdom, but the physiological drivers of infanticide versus care or neglect are relatively unexplored. Here, we identified salient environmental and physiological antecedents of infanticide in the mimic poison frog (*Ranitomeya imitator*), a biparental amphibian in which female parents feed their tadpoles unfertilized eggs. Specifically, we explored potential environmental cues influencing infant-directed behavior by evaluating changes in the frequency of food provisioning and tadpole mortality after either cross-fostering tadpoles between family units or displacing tadpoles within the terraria of their parents. We found that changes in offspring location reduce care and increase infanticide. Specifically, parents fed their displaced offspring less and, in some instances, tadpole mortality increased. We also investigated whether care and infanticide were related to changes in steroid hormone concentrations in an unfamiliar setting. Infanticide of fertilized eggs and hatchlings in the new territory included cannibalism and was associated with lower testosterone concentrations, but not with changes in corticosterone. Overall, our results support earlier findings that familiarity with offspring location drives parental investment in poison frogs, while indicating an association between low androgen levels and infanticidal behavior in an amphibian.

**Highlights:**

- Offspring location drives parental decisions of care vs. infanticide.
- In novel territories, adults cannibalize conspecific, unrelated young.
- Lower circulating testosterone in novel territory is associated with infanticide.

## 1. Introduction

Animal parents exhibit remarkable variation in behavior toward conspecific young, performing parental care (e.g. feeding, protecting, and transporting young), neglect, and infanticide, regardless of kinship (Hrdy, 1979; Royle et al. 2012; Lukas and Huchard, 2014; Lukas and Huchard, 2019). Care requires significant energy investment, rendering parental decision-making sensitive to evolutionary trade-offs dependent on environmental and physiological constraints (Clutton-Brock, 1991; Alonso-Alvarez and Velando, 2012). Thus, parental behavior is often informed by cues that help discriminate between related and unrelated offspring, including indirect identifiers such as location (Stynoski, 2009; Huang and Pike, 2011; Ringler et al., 2016, 2017) and more direct sensory signals like acoustics and smell (Neff and Sherman, 2005; Vergne et al., 2011). While behavioral ecologists have reported numerous adaptive rationales for infanticide across diverse animal groups (Hrdy, 1979; Ebensperger and Blumstein, 2007; Bose, 2022), evidence linking infanticide to hormones is limited outside of mammals. Understanding the physiological mechanisms and evolution of parental responses to offspring requires connecting infanticide to physiological states in a broad range of animal species.

Steroid hormones present a compelling avenue to investigate the physiological basis of infanticide. Infanticide may be considered a form of social aggression, which is often studied in the context of elevated androgens, like testosterone, and elevated glucocorticoids, like corticosterone, especially in males (Summers et al., 2005; Haller, 2014; Wingfield et al., 1990; Hirschenhauser and Oliveira, 2006; Rodríguez et al., 2022). Changes in steroid hormone concentrations that mediate socially aggressive responses can persist even after the behavior occurs (Rodríguez et al., 2022). Within female vertebrates, previous authors have proposed a link between testosterone, parental care, and aggression in mammals, birds, lizards, and fishes (Rosvall, 2020), but also see Bentz et al. (2019), where the relationship between testosterone and aggression in females is less clear. Investigations in both mammalian and non-mammalian taxa indicate that social aggression may be facilitated by the acute elevation of peripheral glucocorticoids (Summers et al., 2005; Haller, 2014). Still, the lack of studies clarifying the physiology of infanticide in non-mammalian taxa prevents the development of a general, comparative, and predictive framework for decision-making toward young.

Poison frogs display an extraordinary diversity of parental care strategies, including tadpole transport, maternal egg provisioning, and cannibalism (Weygoldt, 1987; Summers and Tumulty, 2014). Previous work in poison frogs has suggested that offspring location, territorial status, and recent reproductive activity can influence decisions to perform care or infanticide (Stynoski, 2009; Ringler et al., 2016, 2017; Spring et al., 2019). For example, territorial status determines the performance of tadpole transport or cannibalism in brilliant-thighed poison frog (*Allobates femoralis*) males, while females transport tadpoles based on recent reproductive activity and the exact location of offspring within their territory (Ringler et al., 2016, 2017). These findings have established a framework for investigating environmental cues for infanticide in other poison frogs, but the hormonal correlates of this behavior remain unexplored.

We used mimic poison frogs (*Ranitomeya imitator*) to better understand how external cues elicit hormonal shifts and infanticide. Mimic poison frogs are biparental and sexually monogamous, forming pair bonds that last several months and performing sex-specific parental roles to cooperate in the care of their eggs and tadpoles (Brown et al., 2008). Males of this species transport tadpoles after egg hatching and call females to tadpole sites, whereas females provision trophic eggs to their tadpoles. Additionally, *R. imitator* differ in coloration and patterning based on geographic distribution, and it is unclear whether morph identity may also play a role in offspring recognition.

We used two morphs of *R. imitator (Yurimaguas* and *Rapidos)* to identify key components of offspring recognition, then analyzed adult hormone concentrations during an environmental challenge. We began by identifying salient recognition cues so that we might later expose frogs to a challenge in which they were unlikely to misidentify unrelated young as their own. Based on prior work in poison frogs (Stynoski, 2009; Ringler et al., 2016; Ringler et al., 2017), we hypothesized that offspring recognition depends more on tadpole location than direct recognition cues, including morph-specific features. We performed kin recognition assays, namely cross-fostering within morphs, cross-fostering between morphs, and displacement within parents’ terraria. After measuring egg provisioning and tadpole mortality as proxies for parental investment, we concluded that offspring location is a critical indirect recognition cue for this species.

We then leveraged this finding to conduct simulated territory takeover trials, where adults were individually moved to an unfamiliar environment and simultaneously exposed to unrelated, conspecific offspring. Our hypothesis is that changes in the concentrations of circulating testosterone and corticosterone correlate with infanticidal behavior, which would be reflected by changes in hormone concentrations from baseline to immediately after infanticide. However, it is important to consider that correlations between hormone levels and behaviors do not indicate whether hormone levels drive behavior, whether behavior influences hormone levels, or if another factor is responsible for both Nonetheless, we can glean information on the potentially bi-directional relationship between hormones and behavior using this paradigm. Based on earlier findings in other vertebrates, we anticipated that all adults would exhibit elevated testosterone and corticosterone during this challenge. However, we also predicted that infanticidal adults would have higher testosterone and corticosterone concentrations compared to non-infanticidal adults. We reasoned that testosterone and corticosterone should be reflective of perceived social competition and stress, respectively, and that individuals would react to these challenges by eliminating unrelated young.

## 2. Methods

### 2.1 Animal husbandry

All *R. imitator* eggs and tadpoles in this study were captive-bred in our colony. Adults were purchased from Indoor Ecosystems between 2019-2021 (Whitehouse, Ohio) or Ruffing’s Ranitomeya between 2021-2023 (Tiffin, Ohio, USA). Breeding pairs were housed in 30 x 30 x 45 cm terraria containing sphagnum moss substrate, driftwood, live plants, egg deposition sites, and film canisters filled with water for tadpole deposition. Terraria were automatically misted ten times daily and frogs were fed live *Drosophila* fruit flies dusted with Repashy Calcium Plus (Oceanside, CA, USA) three times per week and springtails once a week. All procedures in this study were approved by the Stanford University Animal Care and Use Committee (Protocol #34242).

### 2.2 Offspring recognition assays

Tadpoles used in this study were observed in home or foster terraria as described (all with the same dimensions and substrates). *Ranitomeya imitator* pairs used for this study were required to have previously successfully raised a tadpole within 31 days of the experiment. Locations of tadpole deposition were marked on the outside of the terrarium with the date and time to ensure consistency in placing tadpole canisters in the same location throughout the duration of the study. Tadpoles were randomly placed into three experimental conditions (**Fig. 1A**): same-morph cross-foster (n = 16 across 6 terraria), different-morph cross-foster (n = 12 across 5 terraria), or intra-tank displacement (n = 18 across 7 terraria). Control tadpoles (n = 26 across 11 terraria) served as a measure of parental investment under normal conditions.

**Figure 1.**
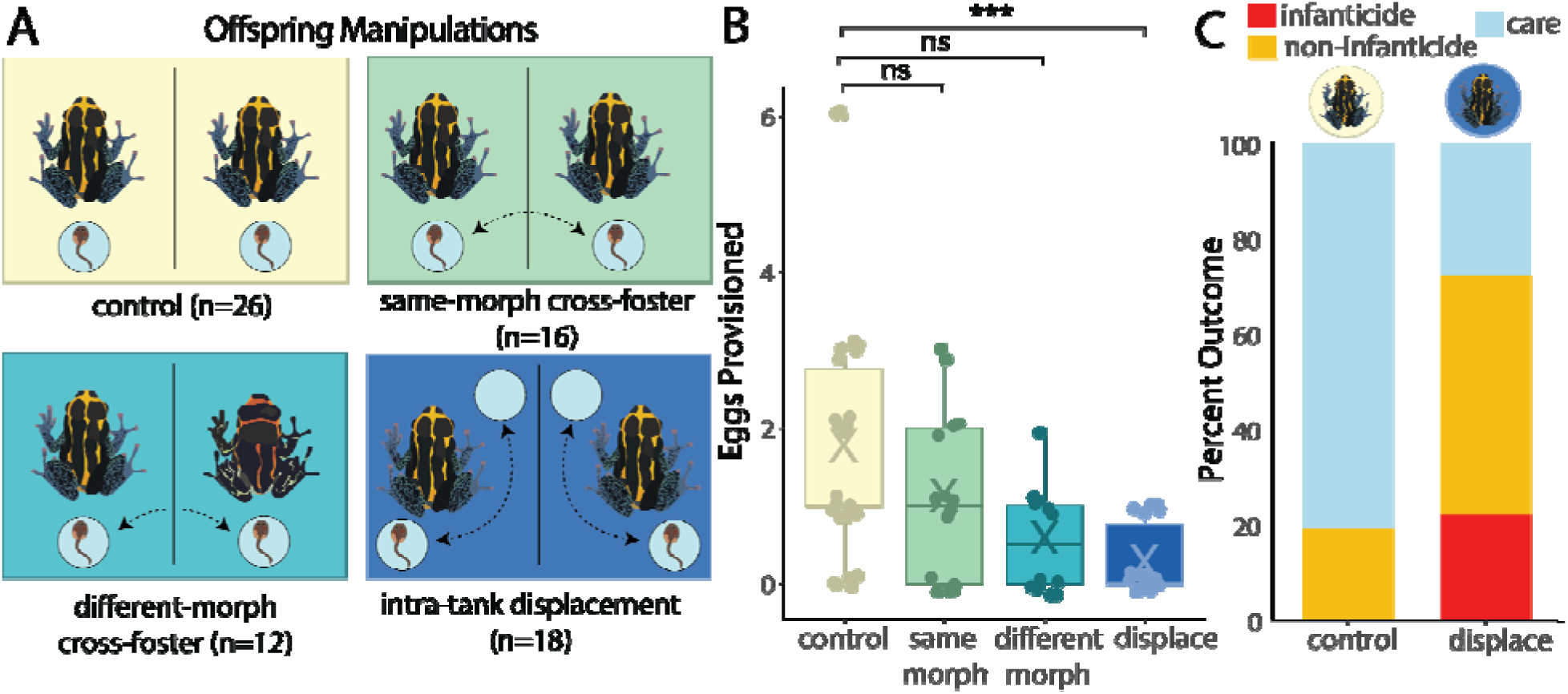
Disrupting offspring location impedes egg provisioning and promotes infanticide. **(A)** Control tadpoles were the offspring of the parents in that terrarium and were not moved from their original locations. Cross-fostered tadpoles were exchanged between terraria, such that they were unrelated to the parents in the new terrarium. They were specifically moved to the same location previously occupied by the offspring they replaced. The different-morph cross-foster manipulation followed the same procedure as same-morph cross-fosters, with the additional condition that tadpoles were exchanged between terraria housing different morphs (striped, banded), discerned by parents’ coloration patterns. Finally, intra-tank displacement tadpoles were related to the parents in the terrarium but were displaced by a minimum of five centimeters to a different location in the terrarium. **(B)** Tadpoles displaced within their parents’ tanks received significantly less feeding in the form of trophic eggs from parents (Tukey adjustment for a family of 4 estimates: p < 0.0008). Significance: 0 < *** < 0.001 < ** < 0.01 < * < 0.05 < ns. “ns” = not significant. **(C)** Only tadpoles displaced within their parents’ tanks (n = 4 out of 18) were subjected to infanticide with no deposition of a second tadpole.

We observed tadpoles in each trial for 14 days and measured trophic egg deposits as a proxy for parental investment. We randomized treatments across tadpoles, using individuals below Gosner stage 32 (Gosner, 1960) to avoid counting feeding patterns that resulted from changes in investments as tadpoles neared metamorphosis. We collected data for every tadpole under controlled conditions before randomly applying treatments. Sampling multiple (one to four) tadpoles per terrarium allowed us to capture focal pair variation in parental care.

Tadpoles were always transported in their respective canisters to retain any tadpole- specific olfactory cues already present in the water and to minimize stress. Tadpoles displaced within their parents’ terraria (“intra-tank displacement”) were moved at a minimum of 5 cm and maximum of 30 cm, depending on availability on the terrarium floor. We purposely selected a minimum distance greater than 2 cm, as this was the minimum distance for indirect offspring discrimination identified in other poison frogs (Stynoski, 2009). Tadpoles were moved either to a laterally opposite corner of the terrarium or elsewhere along the terrarium edge. When undergoing this manipulation, tadpoles were always moved to a location not previously occupied by a canister.

### 2.3 “Takeover” behavioral assays

We designed an assay to identify hormonal changes in infanticidal versus non-infanticidal parents during a simulated territory takeover based on Ringler et al. (2017). We selected paired adults that raised at least one offspring in their terrarium within 31 days of the takeover trial as a proxy for parental experience and reproductive status. Frogs from the same reproductive pair were put into separate takeover trials at the same time to mitigate any risk of mate absence influencing behavior directed toward offspring. The takeover terrarium had the same dimensions, with elements of the environment (vegetation, replacement of dead leaves and substrate, fresh water) disrupted and replaced to introduce unfamiliarity. Rearrangements and cleaning occurred between trials. Takeover terraria always housed at least two tadpole canisters, one upright and filled with water and the second empty and sideways, to remove bias for water availability and shelter, respectively. At the beginning of each trial, frogs were provided with a dish of springtails available *ad libitum* for the length of the trial and were provided with fruit flies on the same schedule as their home tanks.

While baseline hormone collection of the adult occurred, we placed a single, unrelated fertilized late-stage egg to hatchling (Gosner stage 19-22) on a leaf on the terrarium floor. The leaf was positioned below a Wyze Cam v3 to record behavior using previously described methods (Goolsby et al., 2023). Frogs (n = 12 females, 12 males) were introduced to the terrarium and recorded for a maximum of seven days. Infanticidal behavior was characterized as either consumption of the fertilized egg or hatchling (**Supplementary Video 1**) or as repetitive physical disturbances to the egg, where the snout repeatedly dug at the jelly-like casing in what we interpreted as an attempt at egg cannibalism impeded by the encasement (**Supplementary Video 2**).

### 2.4 Steroid hormone collection and processing

We collected hormones immediately before displacing frogs to the new tank (“baseline”), 24 ± 3 hours later (“move”), and following infanticide or tadpole transport, or after seven days if no such behavior occurred (“final”), with hormone collections occurring at approximately the same time of day or immediately following infanticide (**Fig. 2A-B**). Three subjects demonstrated infanticide on the day after displacement, so hormone measurements counted for both “move” and “final” sampling events. Baseline collection always occurred between 11:30 AM and 2:30 PM to avoid confounding effects of circadian physiology on hormone concentrations. We ultimately analyzed data from 59 samples across 22 individuals.

**Figure 2.**
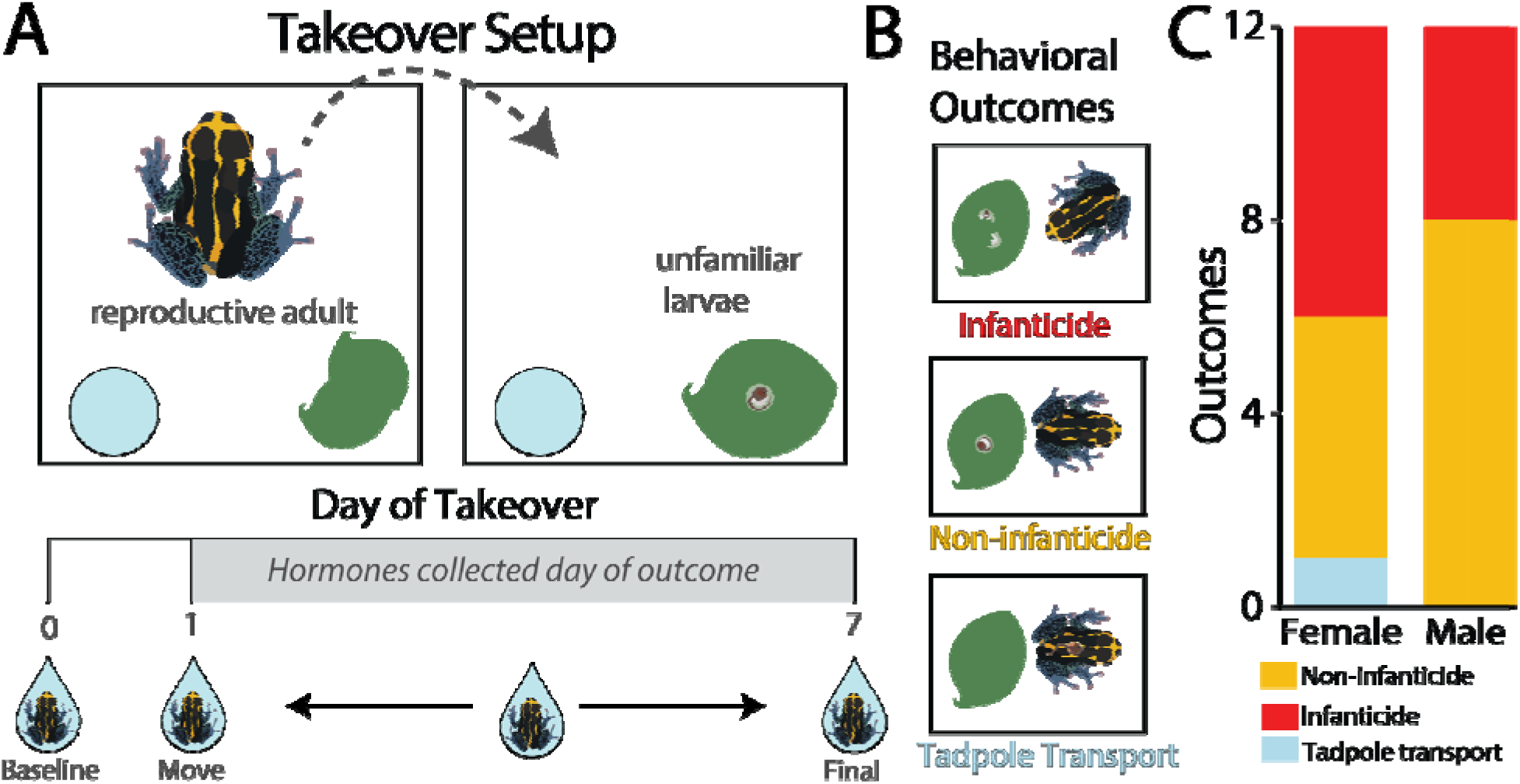
Takeover setup and behavioral outcomes. **(A)** Individual adult frogs were displaced from their home terraria to unfamiliar terraria along with a single, unrelated, fertilized egg or hatchling placed on a leaf. Trials lasted for a maximum of seven days. Hormones were collected via water-borne hormone sampling on the day of the mov (“baseline”), ∼24 hours afterward (“move”), and upon observation of infanticidal behavior, or on Day 7 if no suc behavior occurred (“final”). **(B)** Behavioral outcomes toward eggs or recently hatched tadpoles as observed o motion-trigger Wyze v3 cameras included “infanticide” in the form of cannibalism or physical disturbance to the egg, “transport” when tadpole transport was observed, or “non-infanticide” where these behaviors were not observed. **(C)** Females exhibited all outcomes (non-infanticide: n = 5, infanticide: n = 6, transport: n = 1), unlik males (non-infanticide: n = 8, infanticide: n = 4).

As shown in previous studies with poison frogs and other amphibians (Gabor et al., 2013; Baugh et al., 2018, Baugh and Gray-Gailliard, 2021; Rodríguez et al., 2022; Love et al., 2023), water-borne testosterone and corticosterone were collected as a non-invasive measurement that reflects circulating levels of both hormones. Frogs were individually moved to a petri dish containing 40 mL of distilled water treated with reverse osmosis conditioner to prevent osmotic stress (Josh’s Frogs RO R/x, Osowo, MI, USA). After 60 minutes, we measured body length (snout-vent length) using a digital caliper (Shahe Measuring Tools, Amazon) and body mass using a Maxus precision pocket scale (sensitivity 200 x 0.01 g, Amazon). We also measured mass at each hormone sampling event, then averaged to account for variations related to body fluctuations.

We pre-extracted steroid hormones using Sep-Pak C18 cartridges (Waters, Milford, MA, USA). The cartridges were conditioned using 2 mL of 100% ethanol, followed by 2 mL of Milli-Q water treated with reverse osmosis conditioner. Water samples were pushed through the column at a rate of approx. 10mL/minute, and then eluted with 4 mL of 100% ethanol. Ethanolic extracts were stored in glass vials at 4 □until nitrogen evaporation. One day prior to hormone quantification, the 4 mL of eluted hormone samples were divided into 2 mL new glass tubes for separate corticosterone and testosterone analyses from the same frog. The 2 mL tubes were placed in a 37 water bath and evaporated with gentle flow N_2_ gas. After evaporation, hormone samples were resuspended in 250 μl of assay buffer specific to the corresponding ENZO kit and stored at 4 overnight until analysis the following day.

### 2.5 Steroid hormone quantification via enzyme-linked immunosorbent assays (ELISA)

After overnight incubation at 4 □, samples were warmed to room temperature, and corticosterone and testosterone concentrations were determined using commercially available ELISA kits (ENZO Life Sciences, Farmingdale, NY; Corticosterone: cat. no. ADI-900-097, antibody: donkey anti-sheep IgG, sensitivity: 27.0 pg/mL; Testosterone: cat. no. ADI-900-065, antibody: goat anti-mouse IgG, sensitivity: 5.67 pg/mL) according to the manufacturer’s instructions. We chose to analyze corticosterone rather than cortisol concentrations because corticosterone is suspected to be the main adrenocorticotropic hormone (ACTH)-responsive glucocorticoid for *R. imitator* (Cockrem, 2013; Westrick et al., 2023). Samples were vortexed and 100 µL of the samples were added in duplicate in individual wells of microtiter plates. Following manufacturer instructions, plates were read at 405 nm, with correction between 570 and 590 nm, using a microplate reader (Synergy H1, BioTek Instruments, Winooski, VT, USA).

Hormone concentrations were calculated using a four-parameter logistic curve in the software Gen5 (version 3.05, BioTek Instruments, Winooski, VT, USA). Samples with duplicate measurements that yielded a coefficient of variation (CV) > 30% were excluded from later analyses (corticosterone: 7 of 59 samples, 11.9%; testosterone: 2 of 59 samples, 3.4%). Before removing these duplicates, intra-assay CV values for corticosterone and testosterone were 14.0% and 10.0%. After removal, intra-assay CV values were 8.4% and 9.4%. Inter-assay CV values were 11.8% and 13.7%.

### 2.6 Statistics

We cleaned and analyzed all data using RStudio (R version 4.2.3, R Core Team, Boston, MA). We implemented all statistical analyses using a generalized linear mixed model (GLMM) framework. For each analysis, we used the Akaike Information Criterion corrected for small sample sizes (AICc; Burnham and Anderson, 2004) to choose between alternative model fits. We evaluated standard model diagnostics and tested for outliers and quantile deviations using the ‘*DHARMa’* package (version 0.4.6, Hartig, 2022). All models in this manuscript passed all diagnostic tests. To evaluate the significance of main effects, we performed a Type III analysis of variance with Satterthwaite’s method. We evaluated pairwise comparisons and accounted for multiple comparisons using the Tukey method in the *‘emmeans’* package (version 1.9.0, Lenth, 2023). We estimated effect sizes in multi-termed modeling using the ‘*effectsize’* package (version 0.8.6, Ben-Shachar et al., 2020) and with pairwise effect size comparisons using ‘*emmeans’*. Figures were created using the package ‘*ggplot2*’ (version 3.4.4, Wickham, 2016) and assembled in Adobe Illustrator (Adobe Illustrator 2023).

#### 2.6.1 Offspring manipulations

We analyzed the number of eggs deposited for each tadpole among the four experimental conditions (**Fig. 1A**). To test whether egg deposition changes due to cross-fostering and intra-tank displacement, we used a GLMM with a zero-inflation component implemented using the ‘*glmmTMB*’ package (version 1.1.8, Brooks et al., 2017). We chose a Poisson distribution based on best fit, with eggs fed as a dependent variable, experiment type and number of siblings within a terrarium as independent variables, and focal terrarium as a random effect.

#### 2.6.2 Takeover trials

To determine how testosterone, corticosterone, and testosterone:corticosterone ratios change relative to infanticidal behavior, we implemented a GLMM using the ‘*lmerTest*’ package (3.1-3, Kuznetsova et al., 2017). We set the outcome (“infanticide” vs “non-infanticide”), sampling event (“baseline” vs. “move” vs. “final”), sex, and their interactions as fixed effects and individual frog identity, average individual mass, volume of collection water, and body length (snout-vent length) as random effects. We selected this method rather than linearly normalizing by mass, body length, or water collection volume following correlation analyses (**Figs. S1-2**). We estimated average individual mass as the average body mass across sampling events as this was necessary to account for intra-individual differences in feeding. Since mass was not collected for one individual frog, we used mean imputation to estimate the single missing value as the averaged individual mass across the entire study. To improve interpretation of regression coefficients, we removed insignificant interaction terms using stepwise backward regression.

We ran separate models for testosterone and corticosterone. We were additionally interested in ratios of testosterone to corticosterone, results for which are available in the supplement (**Figs. S3-4**; **Tables S1-3**). We selected appropriate data transformations (natural log, square root, inverse, no transformation) for modeling each dependent variable using AICc. Using AICc and diagnostic criteria, we selected a log transformation for corticosterone and an inverse transformation for testosterone. Ratios were modeled using the untransformed testosterone and corticosterone values to compute an initial ratio value, which was then log-transformed. Because corticosterone, testosterone, and ratio values of samples exceeding the CV threshold of 30% are deemed unreliable, we treated these values as missing data. All of our final models passed all diagnostic tests.

## 3. Results

### 3.1 Offspring location as an indirect recognition cue

To identify which offspring features parents use to recognize and care for their own young, we either cross-fostered or displaced tadpoles within their parents’ terraria and observed egg deposition and tadpole mortality for 14 days. We hypothesized that in addition to offspring manipulations, the number of siblings present may also influence how many eggs can be provisioned. We manipulated offspring identity (same-morph cross-foster, different-morph cross-foster) and offspring location (intra-tank displacement, **Fig. 1A**). We found that offspring manipulation type (^2^ = 17.74, p < 0.0005; **Table S4**) was an overall better predictor of eggs provisioned than the number of siblings present in a tank at a time (CI: -0.51 – 0.22, ^2^ = 0.62, p = 0.43; **Table S4**).

We next asked whether offspring displacement influenced parental provisioning of young and found a strong effect on the number of eggs provisioned (CI: -2.43 – -0.37, z = -2.66, p < 0.01, **Table 1**; model diagnostics may be found in **Fig. S5** with details for analysis of deviance in **Table S4.** Specifically, over 14 days, displaced tadpoles received significantly fewer trophic eggs compared to positive controls (**Fig. 1B**, p_adj_ = 0.0008, = 1.81). Across our experiment, control tadpoles were fed the most, at an average of 1.77 eggs every two weeks, whereas same-morph and different-morph cross-fosters were fed less than positive controls, being fed 1.13 and 0.58 eggs on average, respectively. Displaced tadpoles were the least cared for, being fed an average of 0.27 eggs within two weeks. Tadpoles cross-fostered within and outside of their morph did not experience a significant reduction in feeding compared to positive controls (**Fig. 1B**, p_adj_ = 0.16, 0.55; = -0.409, -0.910; **Table S5**). Tadpoles displaced within their parents’ terraria were subjected to a variety of infant-directed behaviors, including “care” as exemplified by egg feeding, “non-infanticide” as exemplified by the absence of observed feeding, and “infanticide” as shown by physical injury to the tadpole without deposition of a second tadpole, which was only observed in the intra-tank displacement manipulation (**Fig. 1C**).

**Table 1.**
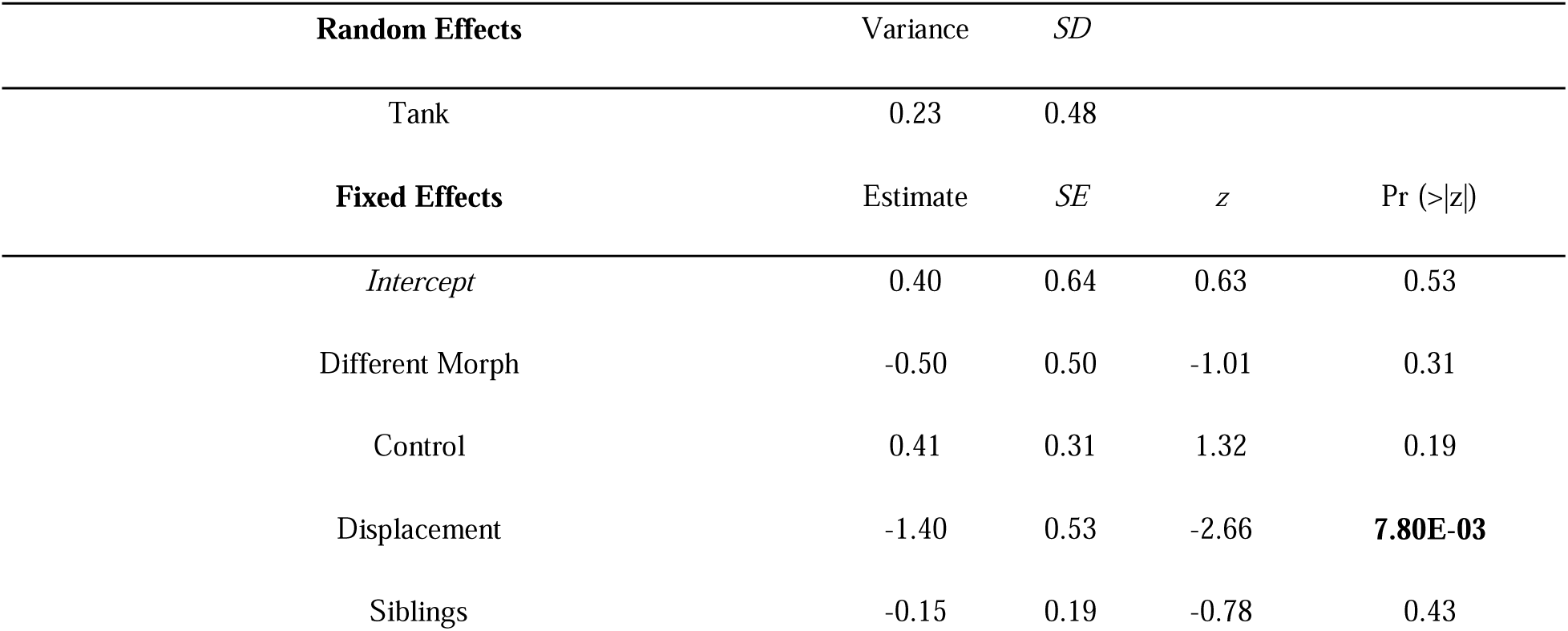
Final model fit of eggs fed with offspring manipulations. SD = Standard Deviation. SE = Standard Error, z = standardized Z value, Pr() = P value.

### 3.2 Infanticide in new territory

After concluding that offspring location is an important factor in parental decisions of care versus infanticide, we performed a “takeover” behavioral assay to identify hormonal correlates of infanticide. We analyzed adult corticosterone and testosterone concentrations with respect to outcome (“infanticide” versus “non-infanticide”), sampling event (“baseline”, “move”, and “final”), and interactions between outcome and event themselves **(Fig. 2A-B**).

Of the twelve trials performed with females, six of the twelve females fell within th “infanticide” outcome, while the other five were categorized as ignoring the conspecific young (“non-infanticide”). One individual demonstrated parental care, performing tadpole transport (**Supplementary Video 3**). As we only observed transport by one individual, we did not have enough statistical power to analyze this result distinct from other non-infanticidal females, and therefore we excluded it from downstream analyses. Of the twelve trials performed with males, eight showed no infanticidal behavior toward conspecific young while four performed infanticide (**Fig. 2C**). Latency to infanticide decisions were variable, with the behavior occurring on a range of days, from the day after movement to the new terrarium to the full seven days (**Fig. S6**). In addition to interactions with offspring, we measured frog mass at each sampling event. The median frog mass decreased from baseline to final sampling (**Fig. S7**).

### 3.3 Infanticide and corticosterone

Log corticosterone (corticosterone hereafter) concentrations did not vary significantly between infanticidal versus non-infanticidal adults (**Table 2**, η_p_² = 0.01, t = -0.59, p = 0.56; **Fig. 4A, Table S6**) nor did they vary significantly between males and females (**Table 2**, η_p_² = 0.17, t = 2.00, p = 0.06; **Fig. 4A, Table S6**). On average, corticosterone concentrations increased approximately 54 pg/mL during takeover of novel territory, although this increase was not significant (η_p_² = 0.12, t = 1.69, p = 0.10, **Table 2**). Model diagnostics may be found in **Fig. S8** with analysis of deviance details in **Table S6**. No pairwise comparisons between combinations of behavioral outcomes, sampling events, and sex were significant (**Table S7**).

**Table 2.** Final corticosterone model fit of log(corticosterone) by sampling event, behavioral outcome, and sex. Frog identity, frog length, and average mass are random effects. SD = Standard Deviation. SE = Standard Error, t = t-statistic, Pr (>|t|) = P value.

| Random Effects | Variance | SD |  |  |  |
| --- | --- | --- | --- | --- | --- |
| Body length | 0.03 | 0.18 |  |  |  |
| Average mass across trial | 0.05 | 0.22 |  |  |  |
| Residual | 0.09 | 0.30 |  |  |  |
| Fixed Effects | Estimate | SE | df | t | Pr ($>\vert t\vert $) |
| *Intercept* | 6.25 | 0.14 | 30.92 | 44.04 | **<2.00E-16** |
| Outcome | -0.08 | 0.14 | 23.15 | -0.59 | 0.56 |
| Move | 0.19 | 0.11 | 27.63 | 1.69 | 0.10 |
| Final | -0.04 | 0.11 | 36.94 | -0.32 | 0.76 |
| Sex | 0.30 | 0.15 | 20.18 | 2.00 | 0.06 |

### 3.4 Infanticide and testosterone

Across all samples, the inverse transformation of testosterone (testosterone hereafter) concentration varied significantly with sampling events (η_p_² = 0.31, p=0.03, F=4.12, **Fig. 3B, Table S8**). Testosterone concentrations also changed between behavioral outcomes (η² = 0.28, p=0.02, F = 6.40, **Fig. 3B, Table S8**), but were not sex-specific (η_p_² = 0.04, p=0.40, F=0.75, **Fig. 3B, Table S8**). Infanticidal individuals, across their trial, averaged about two-thirds of the testosterone concentrations (67.94 pg/mL) of non-infanticidal individuals (97.81 pg/mL). In other words, testosterone decreased, on average, during potentially stressful events (baseline compared to move), and during infanticidal behaviors. Most notably, testosterone concentrations significantly decreased 24 hours after takeover onset (t = 3.30, p < 0.005), but the extent of this decrease was significantly dependent on whether the individual eventually performed infanticide (η_p_² = 0.35, t = -2.60, p = 0.02, **Table 3**). Model diagnostics may be found in **Fig. S9** with full analysis of deviance details in **Table S8.**

**Figure 3.**
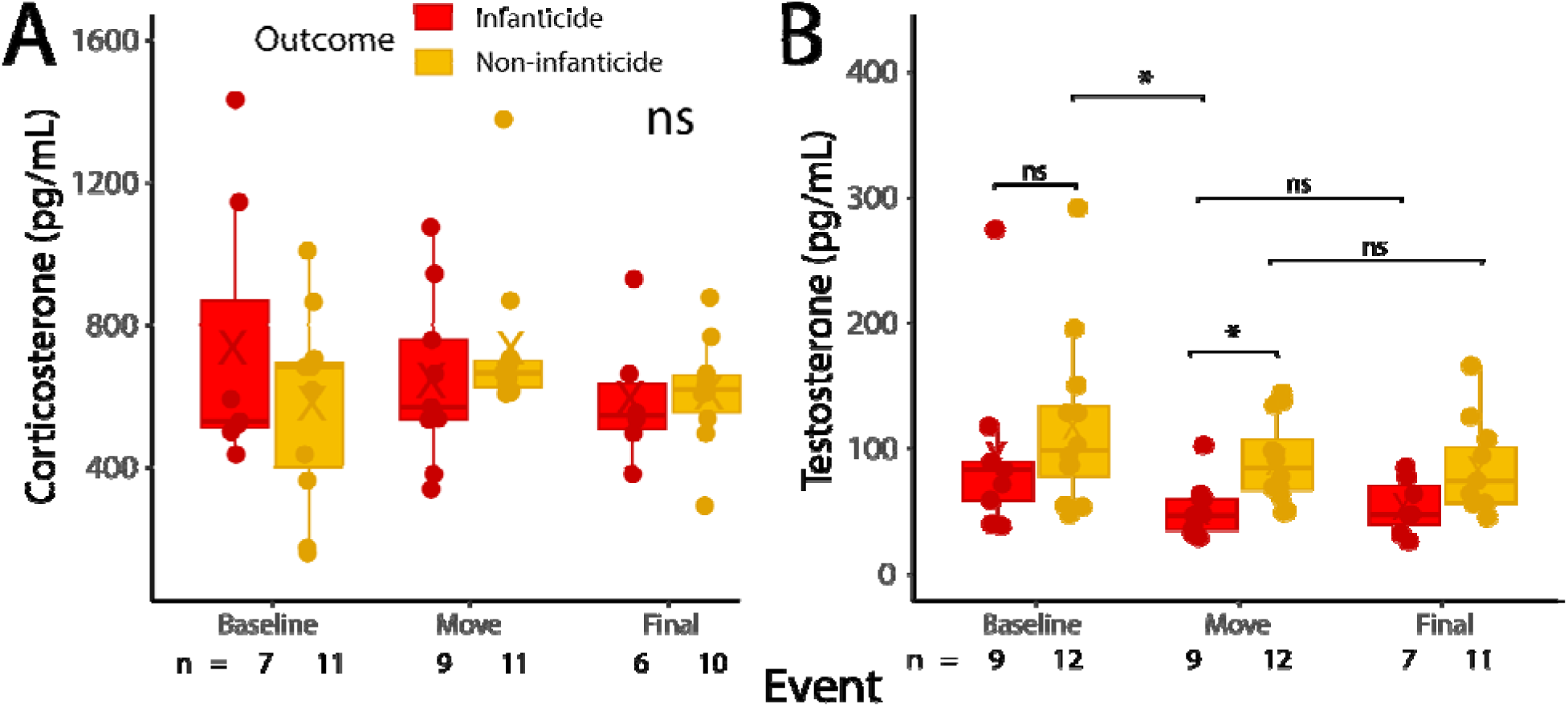
Hormone concentrations by event and behavioral outcome. **(A)** Corticosterone and **(B)** testosteron concentrations by sampling events and behavioral outcomes. The “X” in each box represents the average. Th horizontal lines in each box plot represent the medians of each group. All results are shown in raw units.

**Table 3.**
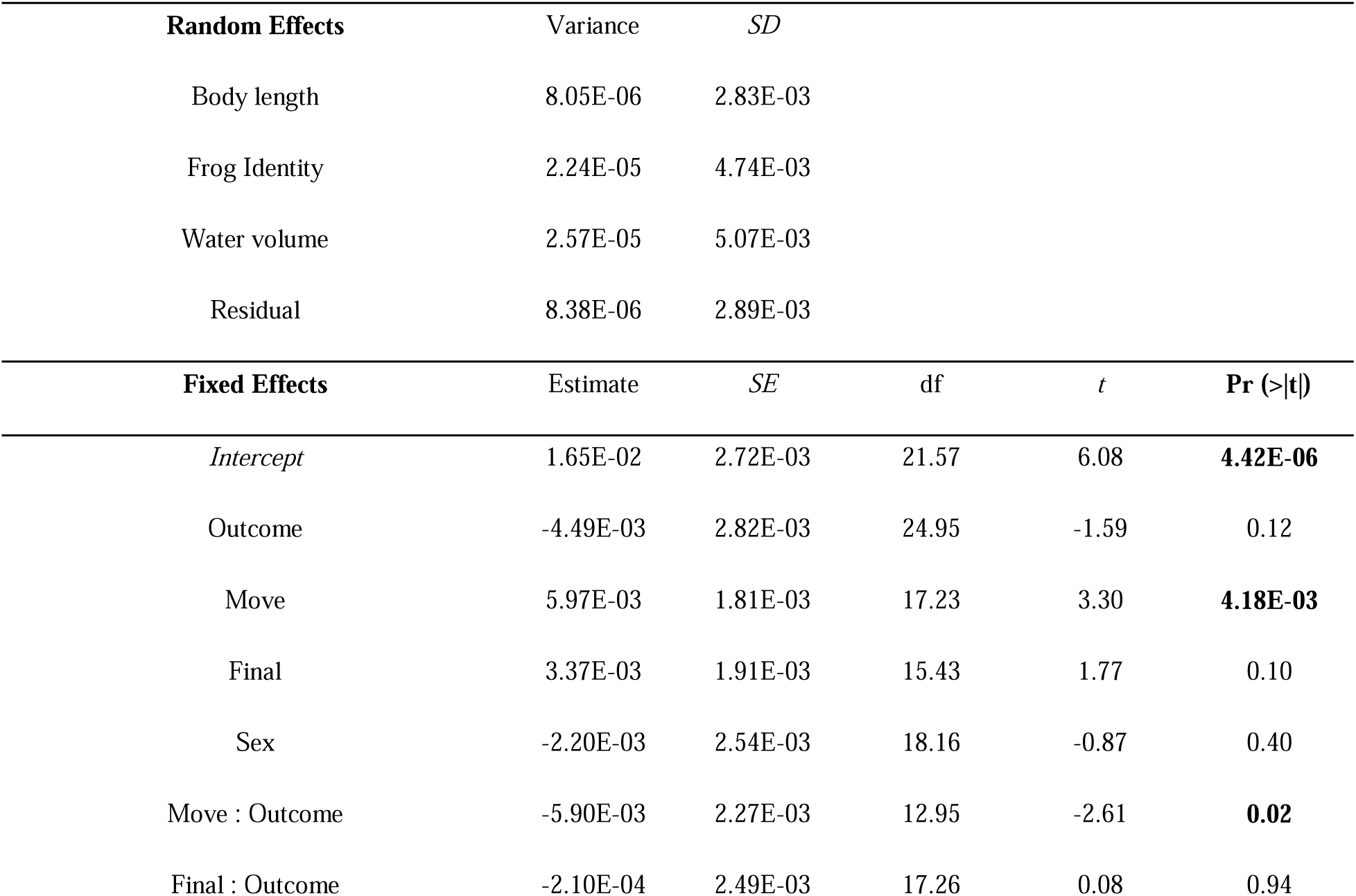
Final testosterone model fit of 1/(testosterone) by sampling event, behavioral outcome, and sex. Frog body length, identity, and water collection volume are random effects. SD = Standard Deviation. SE = Standard Error, t = t-statistic, Pr (>|t|) = P value.

In pairwise comparisons, we examined testosterone concentrations in relation to interactions between sampling events and outcomes. We found that baseline testosterone concentrations in adults who did not perform infanticide were significantly greater than testosterone concentrations in infanticidal individuals moving into a new territory (Cohen’s *d* = -3.62, t ratio = 3.38, p = 0.03). Finally, infanticidal and non-infanticidal individuals have different testosterone concentrations when moved into a new territory (average of 50.62 pg/mL and 90.12 pg/mL, respectively; Cohen’s *d* = 3.59, t ratio = 3.38, p = 0.03). Within infanticidal individuals, we found a moderately positive (r = 0.48), albeit statistically insignificant, correlation between testosterone concentrations 24 hours into the territory takeover assay and latency to infanticide (p = 0.19; **Fig. S10**).

## 4. Discussion

In this investigation, we first delineated tadpole-based features that influence parental decisions to perform care or infanticide, identifying location as a salient indirect kin recognition cue. Then, we documented adult behavior toward eggs and hatchlings in unfamiliar territories, recording infanticide for the first time in *Ranitomeya* frogs. By analyzing concentrations of water-borne corticosterone and testosterone, we demonstrated that lower testosterone concentrations precede infanticide in this species.

### 4.1 Parental decision-making in familiar and unfamiliar territories

We observed that displaced tadpoles experienced significant decreases in trophic egg deposition compared to control tadpoles, unlike tadpoles cross-fostered within or between morphs **(Fig. 1B)**. Like other poison frogs (Stynoski, 2009; Ringler et al., 2016; Ringler et al., 2017), *R. imitator* parents likely discriminate between young based on location. Offspring location therefore appears to be a critical cue driving parental care across poison frogs despite diversity in care systems and ecological constraints. Critical decisions between care or cannibalism depending on internal physiological states are especially abundant in anamniotes and squamates (Ray and Maruska, 2023). For example, spawning appears to impede cannibalism in parenting African *Neolamprologus caudopunctatus* cichlids, resulting in the care of foreign young (Cunha-Saraiva et al., 2018). Similarly, brooding children’s pythons (*Antaresia children*) will provide care when eggs are cross-fostered or even replaced with stones (Brashears and DeNardo, 2012).

A proportion of displaced tadpoles were found dead shortly after being moved to a new location in their home terrarium, which we interpreted to be caused by infanticide (**Fig. 1C**). These offspring did not encounter any other tadpoles, making intra-sibling aggression unlikely, despite being well-documented in *Ranitomeya* (Brown et al., 2009; Schulte and Mayer, 2017; McKinney et al., 2022). Offspring cannibalism as a behavioral consequence of intraspecific competition has been previously suggested in other poison frogs (Summers, 1989), where adults might commit infanticide of unrecognized offspring to prevent misdirected care, to promote survival of their own offspring by parasitizing occupied deposition sites, or to prevent offspring cannibalism by unrelated tadpoles. Other species of *Ranitomeya* have been found to parasitize the care of other parents, indicating brood parasitism as a weapon between conspecific competitors (Poelman and Dicke, 2007; Brown et al., 2009, respectively). To understand why tadpoles were killed rather than eaten by adults, we theorize that dead tadpoles may yield a precious protein source for cannibalistic *R. imitator* tadpoles that adults may deposit. Protein can otherwise be difficult to obtain in low-resource environments and may minimize future costs of females having to provision unfertilized egg meals to nutritionally needy tadpoles (Yoshioka et al., 2016). Thus, we hypothesize that infanticide by *R. imitator* adults evolved for parents to maximize their individual reproductive success and opportunities while minimizing those of potential competitors, following evaluation of their environments.

Following the identification of offspring location as an indirect offspring recognition cue, we analyzed the behavioral responses of reproductive adults to unrelated, conspecific young in a novel terrarium, where adults presumably would not mistake young as their own. We found that both males and females cannibalized the stimulus, marking the first documented report of adults cannibalizing young in the *Ranitomeya* genus and supporting previous reports of non-filial cannibalism in other poison frogs (Townsend et al., 1984; Summers, 1989; Ringler et al., 2017; Spring et al., 2019; Dugas et al., 2023). We considered multiple adaptive rationales for non-filial cannibalism in *R. imitator.* First, cannibalism could be a behavioral decision relevant to a trade-off between acquiring caloric content (which risks eating offspring) and providing care (which risks caring for unrelated young), an idea which has been tested extensively in fish (Bose, 2022). Fertilized conspecific eggs constitute part of other frog diets (Beard, 2007). Furthermore, feeding behavior appears mechanistically linked to parental status across taxa including mammals, birds, and fish (O’Rourke and Renn, 2015; Fischer and O’Connell, 2017). Thus, we reasoned that some frogs might be more likely to cannibalize the stimulus egg or hatchling based on differences in how recently they parented, although all subjects produced a tadpole within the previous 31 days. To account for various degrees of hunger and potential feeding-related effects of parental status, frogs were supplied with food *ad libitum*, so it is unlikely that the frogs in the present study cannibalize to compensate for nutritional deficits. Therefore, it appears more likely that cannibalism occurred as a response to stress or intraspecific competition rather than as a means of acquiring food. However, it may be worthwhile for future investigations to clarify the importance of environmental factors not directly addressed here, including mate access, recent reproductive activity, and territoriality.

### 4.2 Hormonal correlates of infanticide

To clarify whether steroid hormones correlate with infanticide, we analyzed infanticidal behavior and sampling events in relation to corticosterone and testosterone concentrations. We expected to find that adults which performed infanticide would exhibit greater corticosterone concentrations upon displacement to new territory. Instead, corticosterone concentrations were not significantly associated with infanticide at any sampling event (**Fig. 3A**). We were also surprised to find no significant differences in corticosterone between sampling events. However, it is possible that sampling at a time point sooner after displacement would have better reflected the physiological changes associated with an acutely stressful environmental change. Interestingly, glucocorticoids have been proposed to promote both avoidant and approaching behaviors (Terburg et al., 2009), which may relate to decisions to cannibalize and ignore unrelated offspring, respectively. Based on our observations, we conclude that behavioral reactions to unrelated young in this species likely depend more on non-corticosteroid pathways, potentially involving other steroids or neuromodulators.

Circulating androgens are classically associated with territorial aggression in mammals, birds, and recently, other poison frogs (Wingfield et al., 1990; Duque-Wilckens et al., 2019; Rodríguez et al., 2022). Therefore, we expected to find that frogs that performed infanticide upon displacement to a new territory would exhibit greater levels of testosterone. Contrary to what we expected, circulating testosterone appears to be negatively associated with infanticide in this species (**Fig. 3B**). Our finding aligns with investigations in some fish, where plasma 11- ketotestosterone concentrations were lower in cannibals but comparable to those typical of parents (Takegaki et al., 2023). Interestingly, low testosterone is also associated with parental care in male poison frogs (Townsend and Moger, 1987); therefore, it seems plausible that decreased androgens can be associated with both care and infanticide in amphibians. Potential mechanisms for such a relationship may include the local aromatization of testosterone to estradiol, which has been linked to aggression in several mammals, fish, and birds (Trainor et al., 2006; Huffman et al., 2013) and is a conserved process in amphibians (Coumailleau et al., 2015). The co-option of estrogens rather than androgens to facilitate aggression can potentially avoid the costs of elevated testosterone, which can be especially detrimental in parents (Wingfield et al., 2001).

Finally, while our work necessitates hormone collections *after* a behavioral outcome, this does not suggest that hormone collections cause behavioral outputs. Rather, it is equally possible that behavioral outputs may drive variation in hormone levels, which has been richly documented across contexts where animals encounter opportunities such as social ascent, resources, or sexual opportunities (Nelson, 2009; Maruska and Fernald, 2010). We hope that future work can functionally delineate the relationship between androgens and infanticidal behavior in animals broadly.

## 5. Conclusions

In this study, we aimed to clarify environmental and hormonal cues for infanticide in mimic poison frogs. Our results suggest that, consistent with other poison frogs, care and infanticide in this species are antagonistically linked on the basis of a simple external cue: offspring location. We showed that mimic poison frogs perform infanticide in both familiar and unfamiliar territories, wherein they targeted related and unrelated young, respectively. Based on the ecological history of *R. imitator*, we posit that infanticide in this species serves to prevent misdirected care and eliminate intraspecific competition. Although infanticide has been observed in other poison frogs, to our knowledge, this is the first such report in a monogamous or biparental amphibian, indicating that infanticide by both sexes can occur regardless of mating or parenting systems in this clade. Moreover, our hormonal analyses indicate that low concentrations of circulating androgens can precede infanticide following social and environmental perturbations, which contributes a unique perspective to the broader aggression literature. Overall, these findings offer fresh insights into how adults adjust their behavior and physiology to make life-or-death decisions toward offspring in the face of social instability. In the future, a concerted analysis of endocrine and neural activities compared between individuals performing infanticide, neglect, and care is a promising next direction to uncover evolutionary innovations in physiology underlying offspring-directed behaviors.

## CRediT authorship contribution statement

**Amaris R. Lewis:** Conceptualization, Methodology, Formal Analysis, Investigation, Data Curation, Writing - Original Draft, Writing - Review and Editing, Visualization, Project Administration, Funding Acquisition. **Billie C. Goolsby:** Conceptualization, Methodology, Formal Analysis, Investigation, Data Curation, Writing - Original Draft, Writing - Review and Editing, Visualization, Project Administration. **Bryan H. Juarez:** Methodology, Formal Analysis, Resources, Writing - Original Draft, Writing - Review and Editing, Visualization, Supervision. **Madison P. Lacey**: Investigation, Resources, Writing - Review and Editing. **Lauren A. O’Connell**: Resources, Writing- Review & Editing, Funding Acquisition, Supervision.

## Acknowledgments

This research was conducted at Stanford University, which is located on the ancestral and unceded land of the Muwekma Ohlone tribe. We thank Dr. Camilo Rodríguez Lopez (CRL) and Dr. Ricardo Cossio for feedback on this manuscript. We are additionally grateful to CRL for guidance on steroid hormone collection and processing. We also thank the members of the Laboratory for Organismal Biology for providing routine frog care and continuous support. Finally, we kindly thank the two reviewers for their thoughtful comments which improved the quality of this manuscript.

