## Supplementary Figures for "Infanticide is driven by unfamiliarity with offspring location and associated with androgenic shifts in mimic poison frogs"

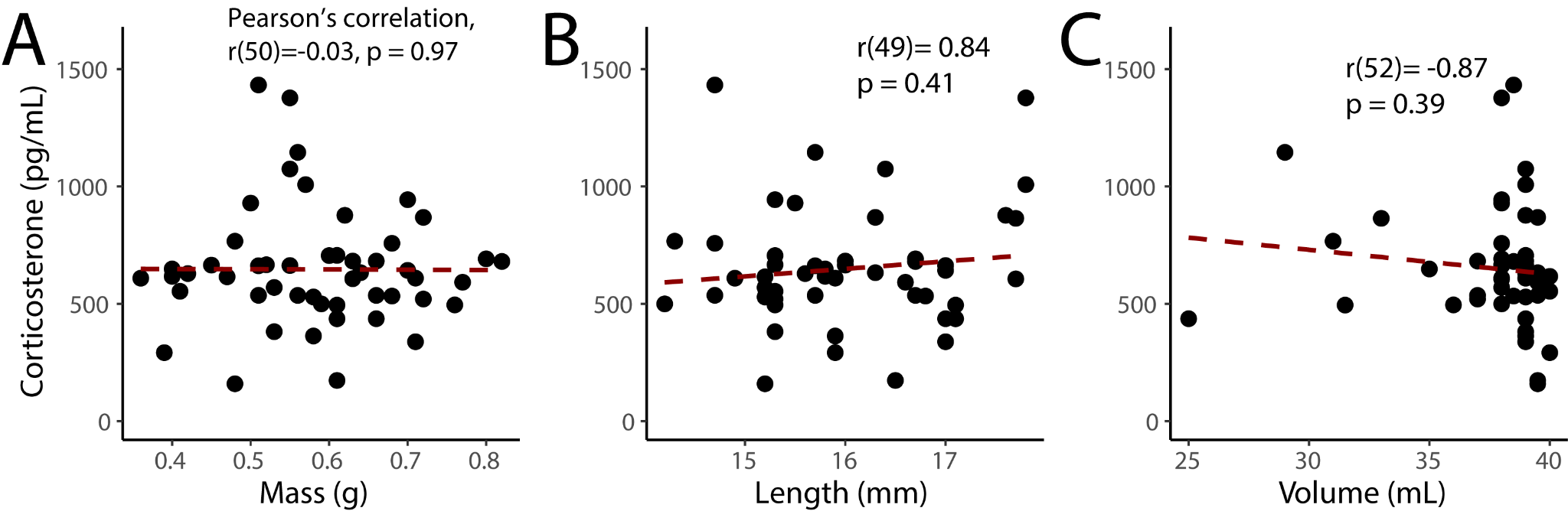
**Figure S1. Distribution of corticosterone values by mass, length, and sample water volume.** Correlations were statistically analyzed by Pearson’s product-moment correlation. (A) No significant correlation was found between mass (g) and corticosterone values (pg/mL); t = -0.22, df = 52, p = 0.83, R = -0.03; (B) No significant correlation was found between length (mm) and corticosterone values (pg/mL); t = 0.55, df = 54, p = 0.58, R = 0.08; (C) No significant correlation was found between water volume (mL) and corticosterone values (pg/mL); t = -0.78, df = 54, p = 0.44, R = -0.11.

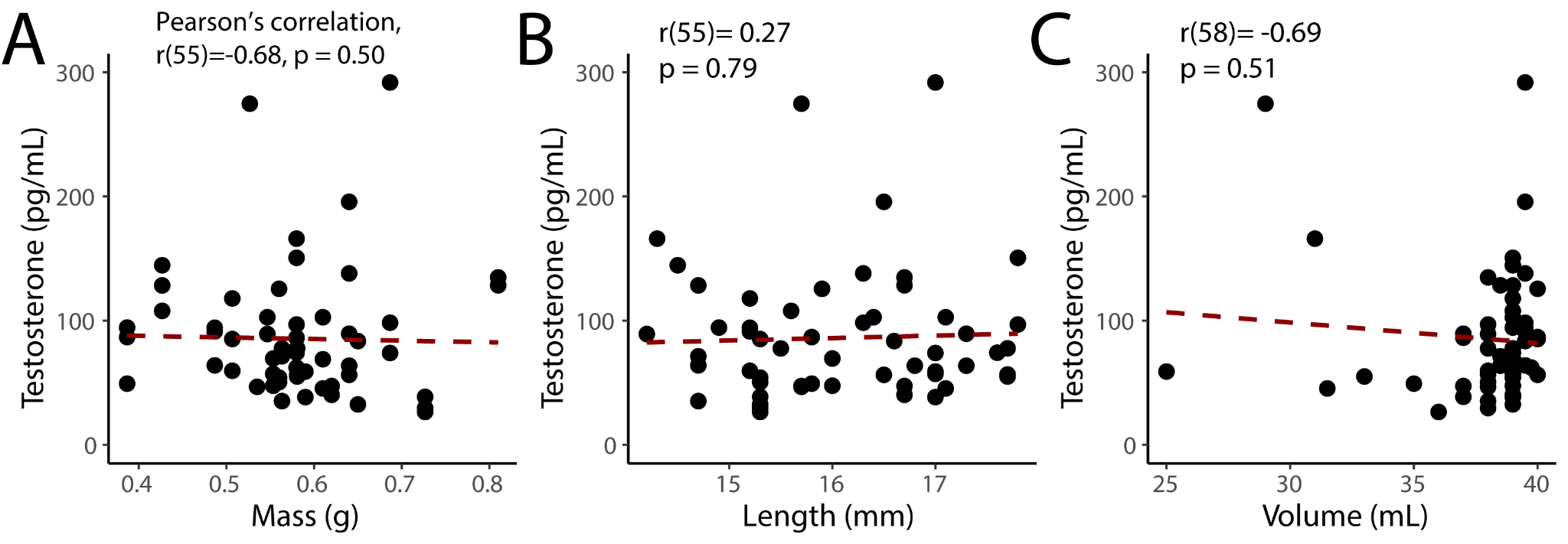

**Figure S2. Distribution of testosterone values by mass, length, and sample water volume.** Correlations were statistically analyzed by Pearson’s product-moment correlation. (A) No significant correlation was found between mass (g) and testosterone values (pg/mL); t = -0.60, df = 55, p = 0.55, R = -0.08; (B) No significant correlation was found between length (mm) and testosterone values (pg/mL); t = 0.32, df = 58, p = 0.75, R = 0.04; (C) No significant correlation was found between water volume (mL) and testosterone values (pg/mL); t = -0.67, df = 58, p = 0.51, R = -0.09.

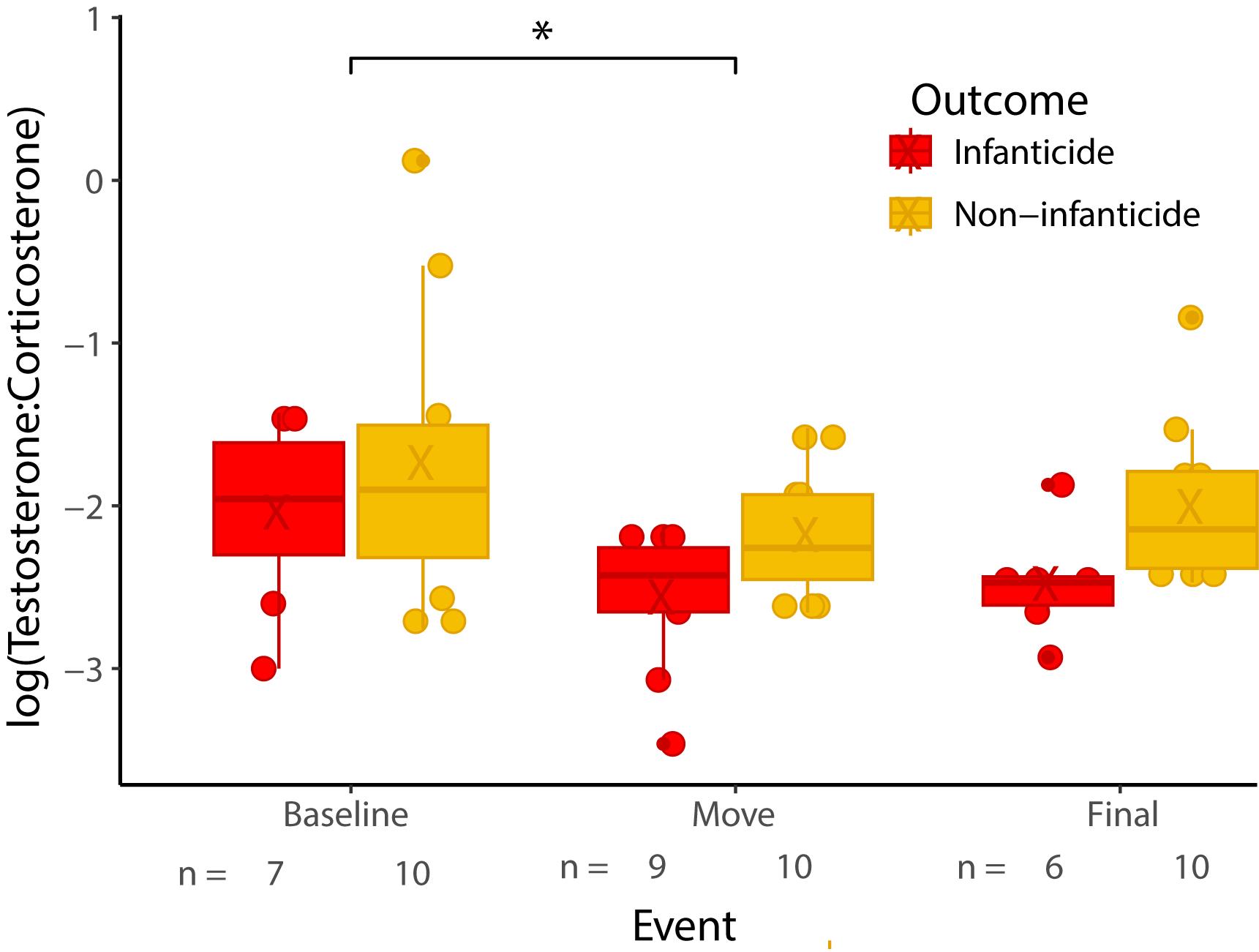

**Figure S3. Natural log of testosterone:corticosterone concentration ratios by sampling events and behavioral outcomes.** Ratios were significantly lower upon displacement to the unfamiliar terrarium. The “X” in each box plot represents the average. The horizontal lines represent the medians of each group. All results are shown in raw units. Signif. codes: 0 < *** < 0.001 < ** < 0.01 < * < 0.05 < . < 0.1 < ns < 1.

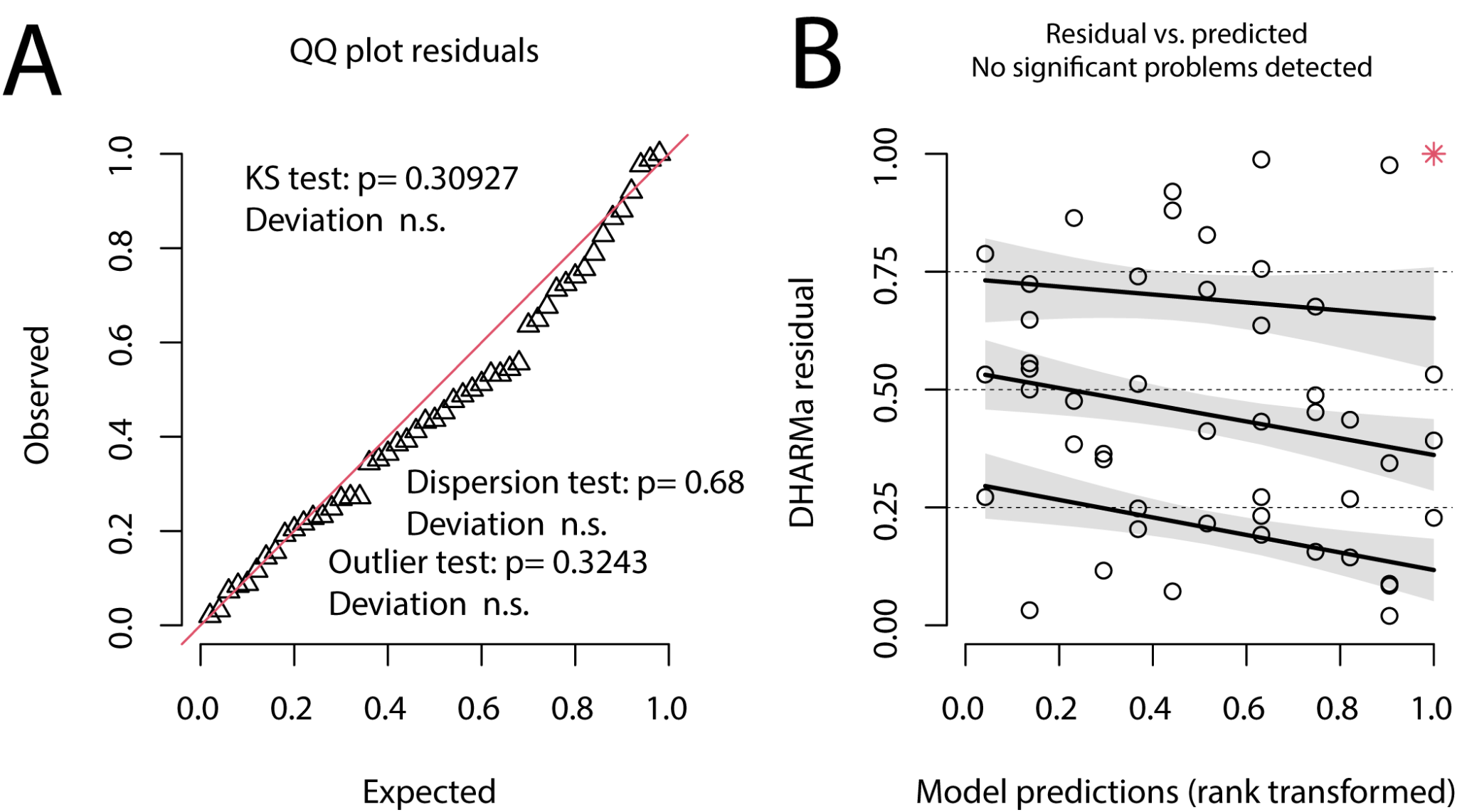

**Figure S4. DHARMa residuals of model fit for ratios.** Residual diagnostics of the following model: mod1.7 <- lmer(log(ratio) ~ outcome + event + sex + (1|length) + (1|id)+(1|avg_mass), data = takeover_hormones_all.TC). Delta AICc of 26. **(A)** QQ plot detects overall deviations from the expected distribution, with tests for correct distribution (Kolmogorov-Smirnov, or KS test), dispersion and outliers. **(B)** Plot of the residuals against the predicted value. Simulation outliers are highlighted as red stars.

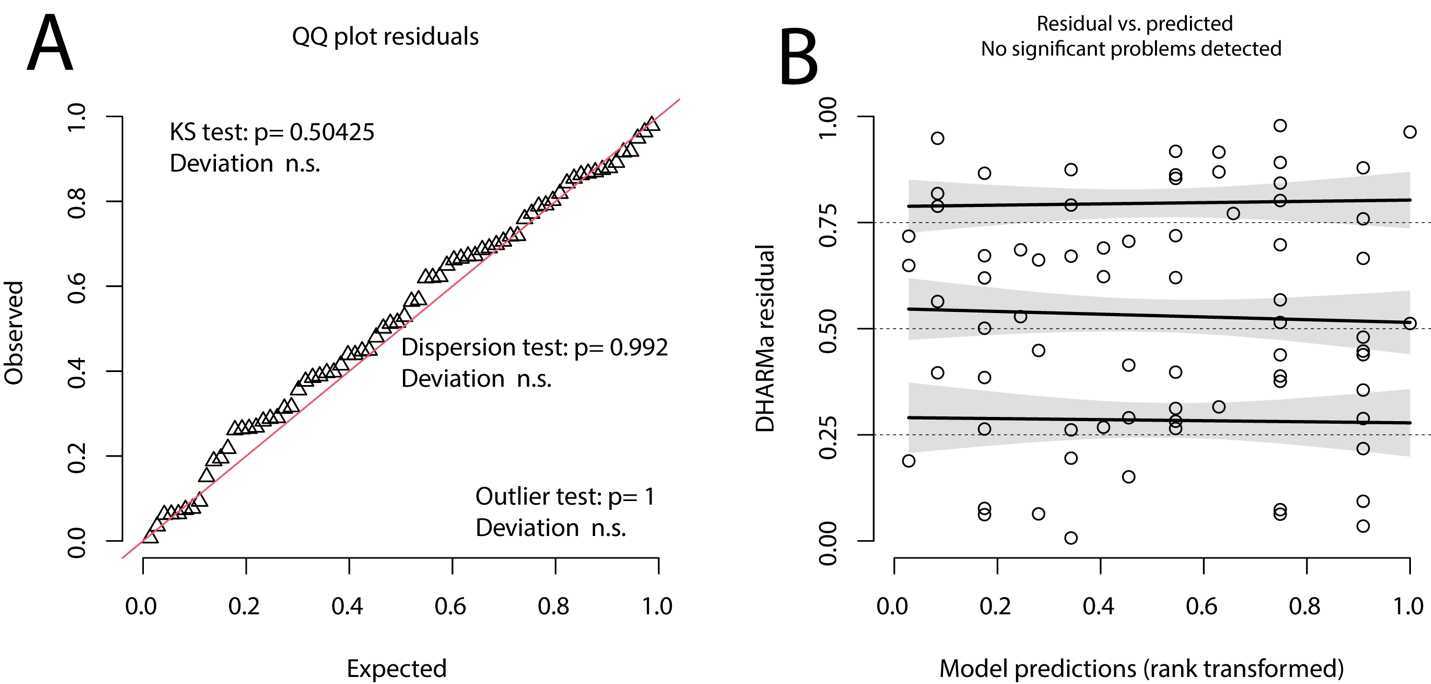

**Figure S5. DHARMa residuals of model fit for offspring manipulations and egg feedings.** Residual diagnostics of the following model: ml3.3 <- glmmTMB(eggs ~ experiment + siblings + (1|tank), family=poisson(), data = eggexperiments). **(A)** QQ plot detects overall deviations from the expected distribution, with tests for correct distribution (Kolmogorov-Smirnov, or KS test), dispersion and outliers. **(B)** Plot of the residuals against the predicted value. Simulation outliers (data points that are outside the range of simulated values) are highlighted as red stars.

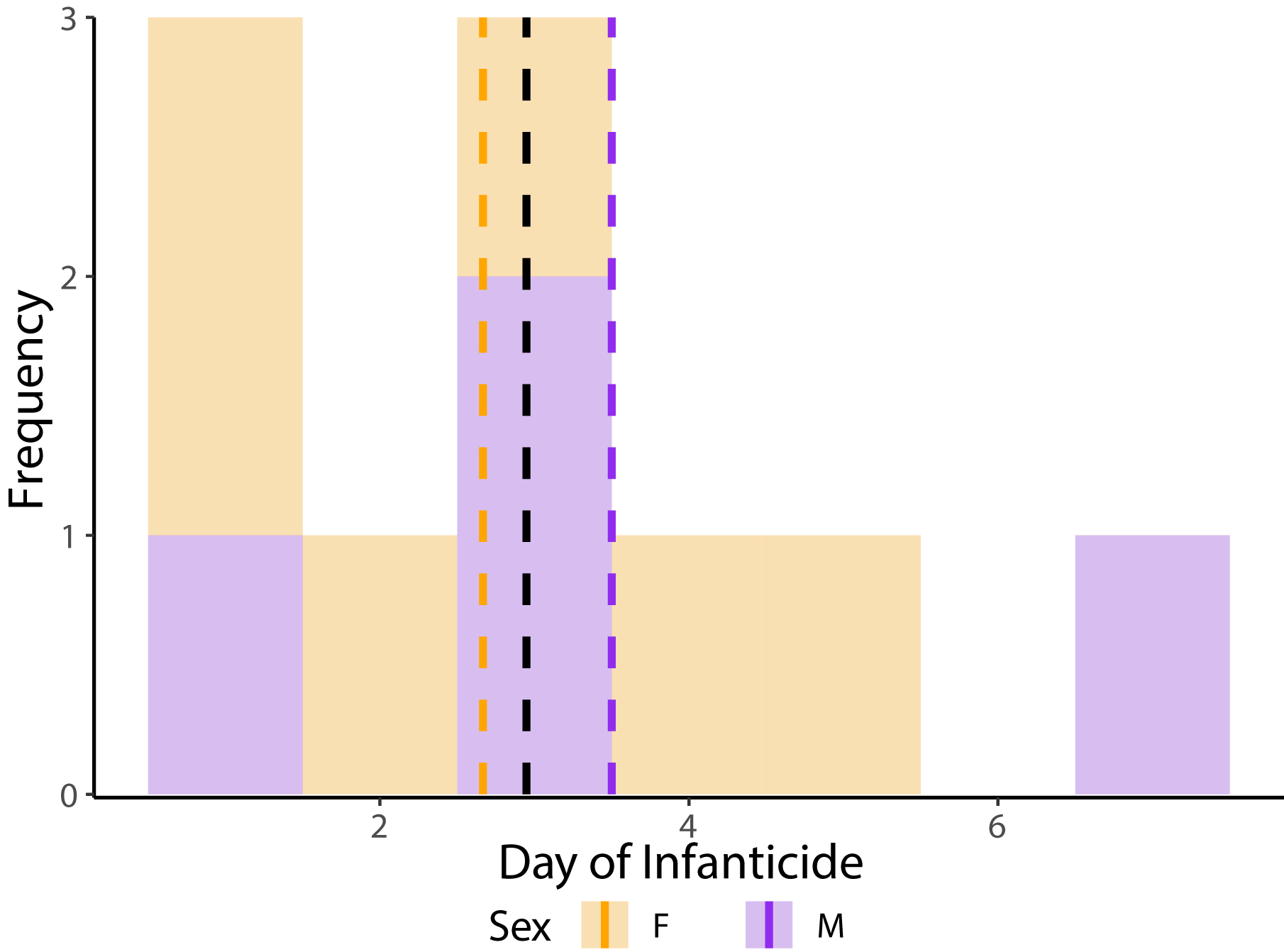

**Figure S6. Distribution of latency to infanticide by sex.** Frequency of males (n = 4) and females (n = 6) that performed infanticide. Orange corresponds with females and purple corresponds to males. Dashed vertical lines represent the average of each sex’s latency to infanticide, with the black vertical line representing the average combining both sexes.

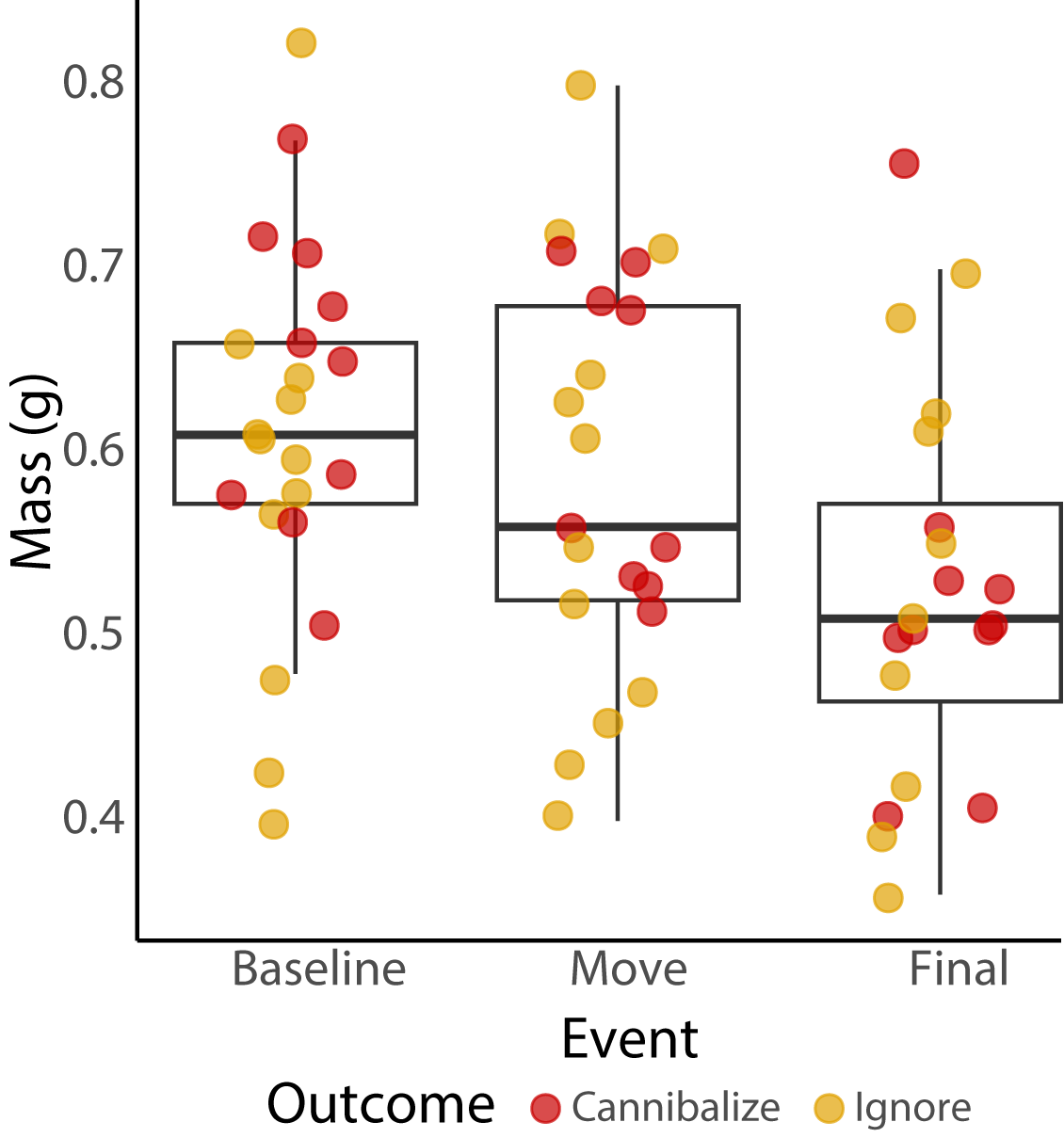

**Figure S7. Frog mass by sampling event.** The median mass of frogs displaced to unfamiliar terraria during takeover trials decreased with time. Median masses at baseline (n = 23), “move” (n = 23), and final (n = 21) sampling events were 0.61 g, 0.565 g, and 0.51 g, respectively.

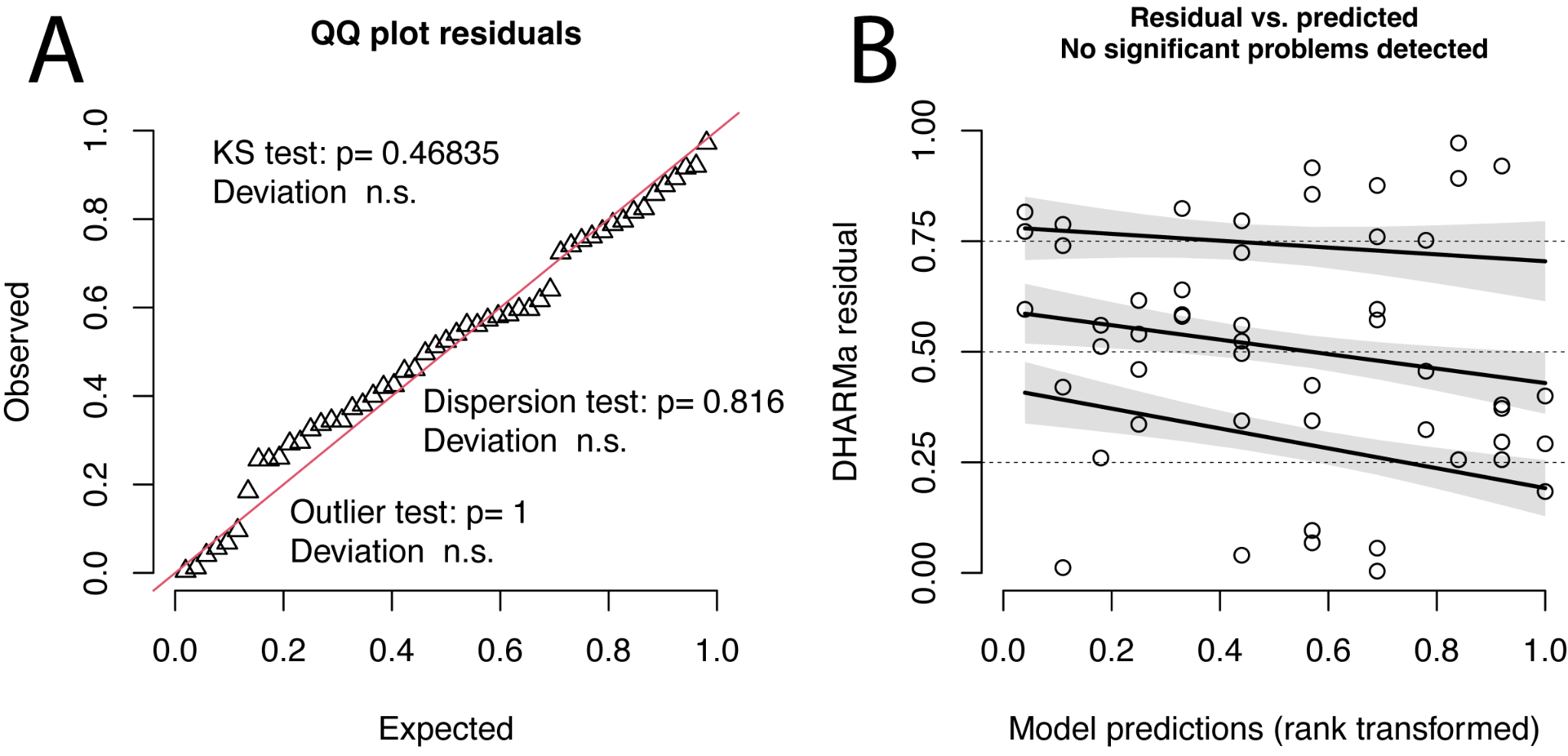

**Figure S8. DHARMa residuals of model fit for corticosterone.** Residual diagnostics of the following model: lmer(log(CORT) ~ outcome + event + sex + (1|avg_mass) + (1|length), data = dat). ΔAICc of 26. **(A)** QQ plot detects overall deviations from the expected distribution, with tests for correct distribution (Kolmogorov-Smirnov, or KS test), dispersion and outliers. **(B)** Plot of the residuals against the predicted value. Simulation outliers are highlighted as red stars.

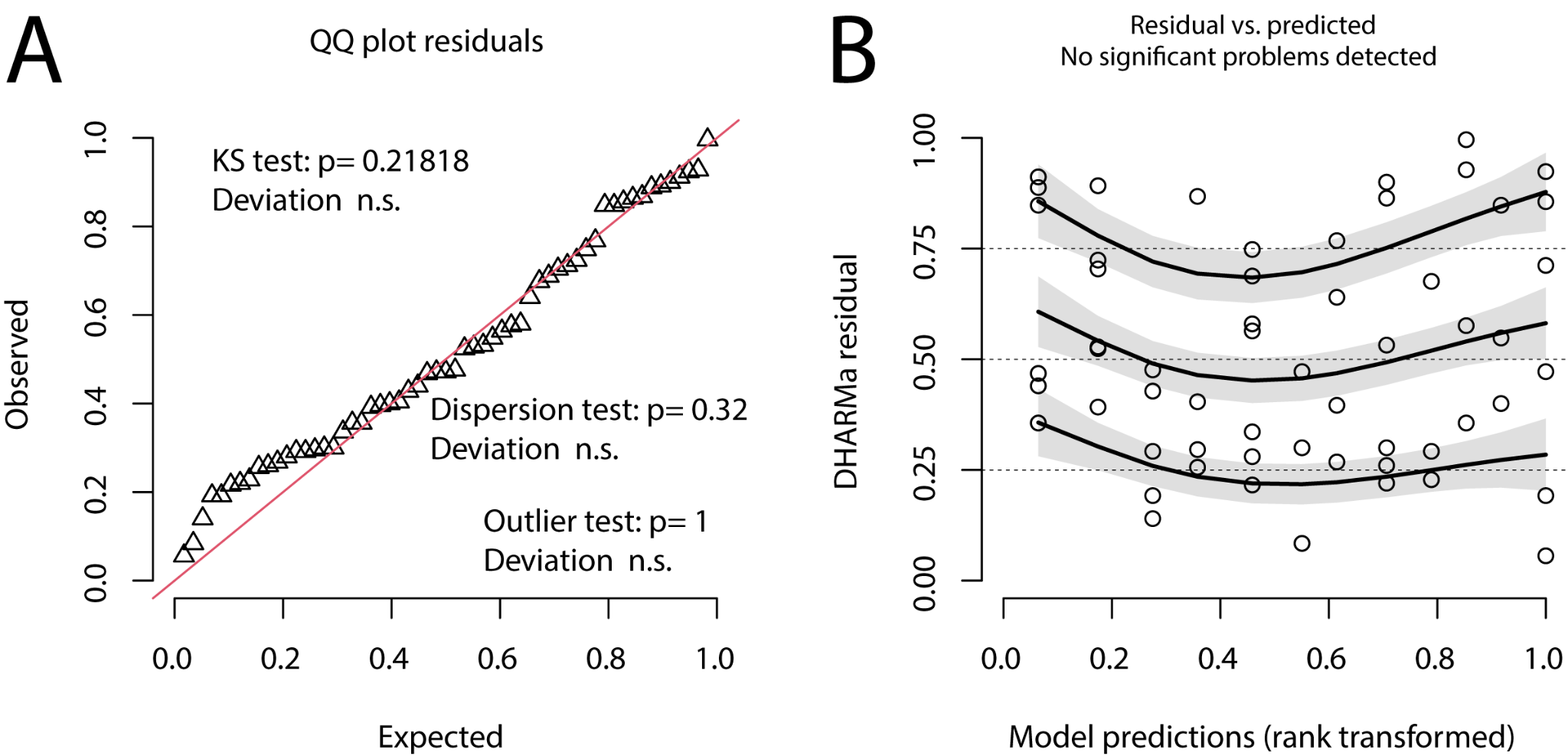

**Figure S9. DHARMa residuals of model fit for testosterone.** Residual diagnostics of the following model: mod3.6 <- lmer((1/(TEST)) ~ outcome * event + sex + (1|id) + (1|volume) + (1|length), data = datT). ΔAICc of 68. **(A)** QQ plot detects overall deviations from the expected distribution, with tests for correct distribution (Kolmogorov-Smirnov, or KS test), dispersion and outliers. **(B)** Plot of the residuals against the predicted value. Simulation outliers are highlighted as red stars.

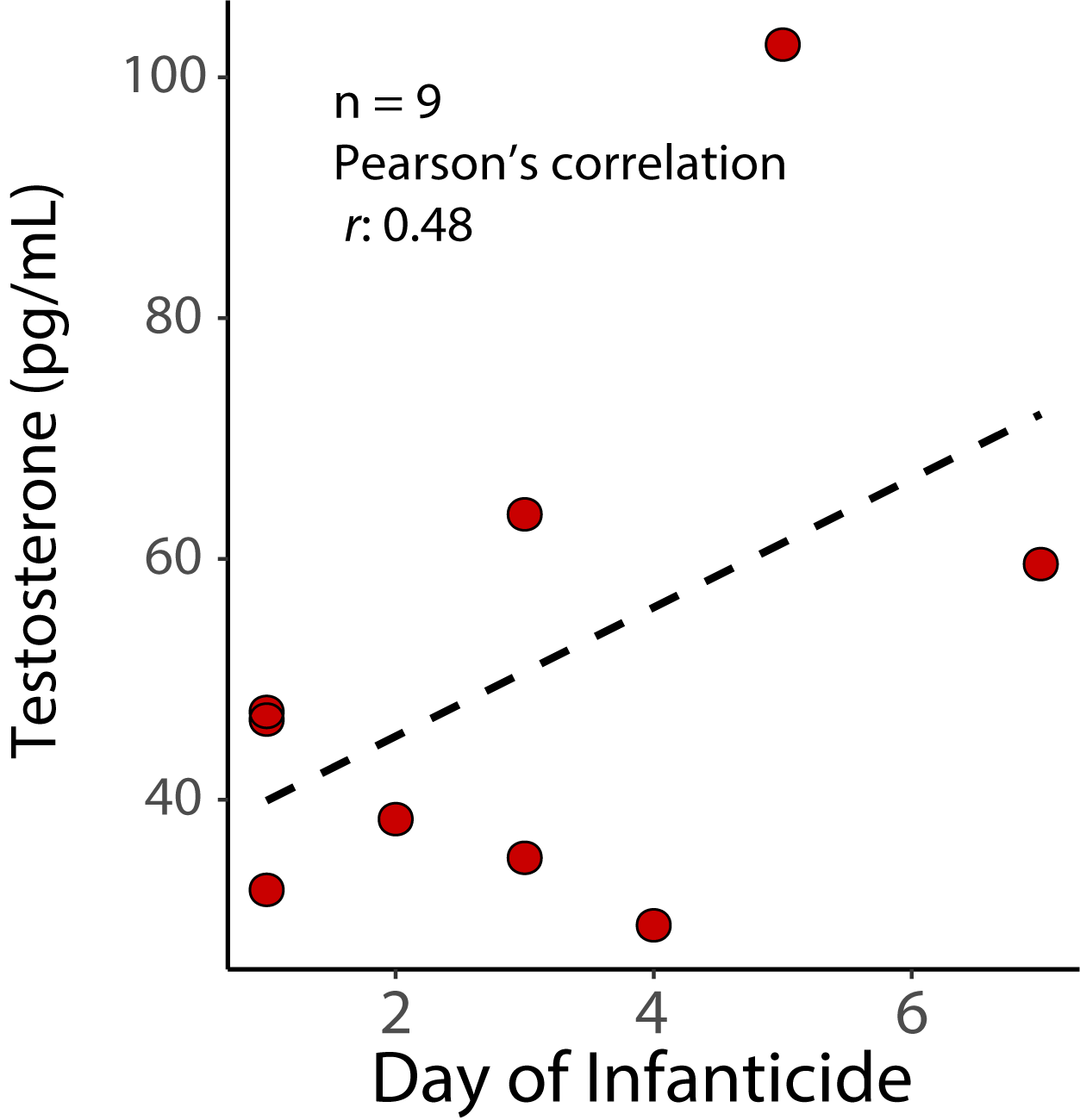

**Figure S10. Testosterone concentrations by day of infanticide.** A moderate, but not significant, correlation between testosterone the day that infanticide was observed (Pearson’s product-moment correlation *r* = 0.48, p = 0.18).
